# Myofibroblasts reduce angiogenesis and vasculogenesis in a vascularized microphysiological model of lung fibrosis

**DOI:** 10.1101/2025.01.10.632378

**Authors:** Elena Cambria, Adriana Blazeski, Eunkyung Clare Ko, Tran Thai, Shania Dantes, David A. Barbie, Sarah E. Shelton, Roger D. Kamm

**Affiliations:** Department of Biological Engineering, Massachusetts Institute of Technology, Cambridge, MA, USA; Department of Mechanical Engineering, Massachusetts Institute of Technology, Cambridge, MA, USA; Center for Excellence in Vascular Biology, Department of Pathology, Brigham and Women’s Hospital and Harvard Medical School, Boston, MA, USA; Department of Medical Oncology, Dana-Farber Cancer Institute, Boston, MA, USA; Joint Department of Biomedical Engineering, University of North Carolina at Chapel Hill and North Carolina State University, Raleigh, NC, USA

## Abstract

Lung fibrosis, characterized by chronic and progressive scarring, has no cure. Hallmarks are the accumulation of myofibroblasts and extracellular matrix, as well as vascular remodeling. The crosstalk between myofibroblasts and vasculature is poorly understood, with conflicting reports on whether angiogenesis and vessel density are increased or decreased in lung fibrosis. We developed a microphysiological system that recapitulates the pathophysiology of lung fibrosis and disentangles myofibroblast-vascular interactions. Lung myofibroblasts maintained their phenotype in 3D without exogenous TGF-β and displayed anti-angiogenic and anti-vasculogenic activities when cultured with endothelial cells in a microfluidic device. These effects, including decreased endothelial sprouting, altered vascular morphology, and increased vascular permeability, were mediated by increased TGF-β1 and reduced VEGF secretion. Pharmacological interventions targeting these cytokines restored vascular morphology and permeability, demonstrating the potential of this model to screen anti-fibrotic drugs. This system provides insights into myofibroblast-vascular crosstalk in lung fibrosis and offers a platform for therapeutic development.

## Introduction

Lung fibrosis refers to a class of diseases characterized by chronic, progressive, and irreversible scarring of lung tissue that disrupts gas exchange (*1*, *2*). While it can result from a variety of conditions, including viral infections (*3*) or exposure to environmental toxins (*4*), a particularly severe form, idiopathic pulmonary fibrosis (IPF), has no known etiology and is associated with a low life expectancy (*5*). While the pathogenesis of IPF is not completely understood, accumulation of myofibroblasts and extracellular matrix deposition are key features of the disease (*6*). Interestingly, vascular remodeling is also implicated in IPF (*7*), but there are conflicting reports on whether vessel density and angiogenesis are increased or decreased in diseased tissue (*8*).

Multiple experimental models have been used to study lung fibrosis without yielding new treatments for IPF, suggesting that the models currently used to study this disease do not adequately mimic IPF pathophysiology. The most common animal model, the bleomycin-treated mouse, has been used to identify hundreds of therapeutic agents, but none have translated to the clinic, perhaps due to the reversibility of the fibrotic phenotype in mice and the inability to recapitulate the chronic and progressive pathology observed in humans (*1*, *9*). Many *in vitro* systems have also modeled fibrosis by differentiating fibroblasts into myofibroblasts, which are contractile cells that express α-smooth muscle actin (α-SMA) in stress fibers, mechanically stabilize tissues after injury, and remodel the extracellular matrix (*10–12*). Myofibroblasts are commonly generated by culturing fibroblasts on two-dimensional (2D) surfaces with transforming growth factor-beta (TGF-β), the main profibrotic signaling cytokine in IPF (*13*), or by growing fibroblasts on pathologically stiff substrates (*14*). However, these simplified models fail to recapitulate the three-dimensional (3D) arrangement, complex biophysical milieu, and heterocellular interactions of lung tissue, limiting their physiological relevance. Thus, there is a need for more physiological experimental models that will improve our understanding of cellular and molecular mechanisms in lung fibrosis and provide platforms for designing and testing therapeutic interventions with better predictive capabilities.

One promising approach is the use of engineered microphysiological systems, 3D platforms that combine multiple, self-organizing cell types with extracellular matrix to recapitulate human tissue microenvironments and architecture (*15*). In particular, engineered vascular networks provide an opportunity to study the interplay between fibrogenesis and vascularization and address its role in IPF pathogenesis. *In vitro* microvascular models of lung fibrosis are rare and have relied on sustained treatment with TGF-β that affect both endothelial cells and fibroblasts (*16*), making it difficult to disentangle the role of myofibroblast-vascular crosstalk in disease progression. In this study, we generated lung myofibroblasts that sustained their phenotype even after withdrawal of TGF-β. We subsequently combined these myofibroblasts with endothelial cells within microfluidic devices to form microvascular models of lung fibrosis. Using this system, we investigated the effects of myofibroblasts on both angiogenesis and vasculogenesis, and identified key cytokines that play a role in mediating these effects. Importantly, we demonstrated the utility of this model for screening anti-fibrotic therapies.

## Results

### Lung fibroblasts treated with TGF-β exhibit a myofibroblast phenotype in 2D

The first step toward creating a microphysiological model of lung fibrosis was activating normal fibroblasts to durably convert them to a myofibroblast phenotype for subsequent incorporation into the 3D culture systems. Human normal lung fibroblasts seeded in 2D (in cell culture flasks or on coverslips) were cultured for 10 days with or without a 1 ng/ml physiological concentration of TGF-β (**Fig. 1**, **Fig. 2A**). The fibroblasts cultured without TGF-β maintained the spindle morphology typical of lung fibroblasts, whereas the fibroblasts exposed to TGF-β changed their morphology to become enlarged and spread, indicating conversion to myofibroblasts (**Fig. 2C**). After 10 days, the expression of key myofibroblasts markers was assessed both at the gene and protein level via RT-qPCR and immunofluorescence, respectively. Compared to the fibroblasts without TGF-β, fibroblasts treated with TGF-β expressed increased levels of ACTA2 (2.7-fold), COL1A1 (7.8-fold), and FN1 genes (2.8-fold), which encode the alpha smooth muscle actin (α-SMA), collagen I, and fibronectin proteins, respectively (**Fig. 2B**). These changes were assessed at the protein level by immunostaining fibroblasts on coverslips and imaging them with a confocal microscope (**Fig. 2C, F, H**). Quantification of the mean fluorescence intensity revealed that fibroblasts treated with TGF-β exhibited a significantly higher actin signal compared to fibroblasts without TGF-β (**Fig. 2D**). While fibroblasts without TGF-β expressed detectable α-SMA protein, fibroblasts treated with TGF-β tended to express more α-SMA. However, this difference was not statistically significant, as measured by mean fluorescence (**Fig. 2E**). Nevertheless, significantly higher fluorescence intensity was observed for collagen I (**Fig. 2F, G**) and fibronectin (**Fig. 2H, I**) in lung fibroblasts treated with TGF-β compared to fibroblasts without TGF-β. These results indicate that treating lung fibroblasts in 2D with 1 ng/ml TGF-β for 10 days induced a myofibroblast phenotype.

**Fig. 1.**
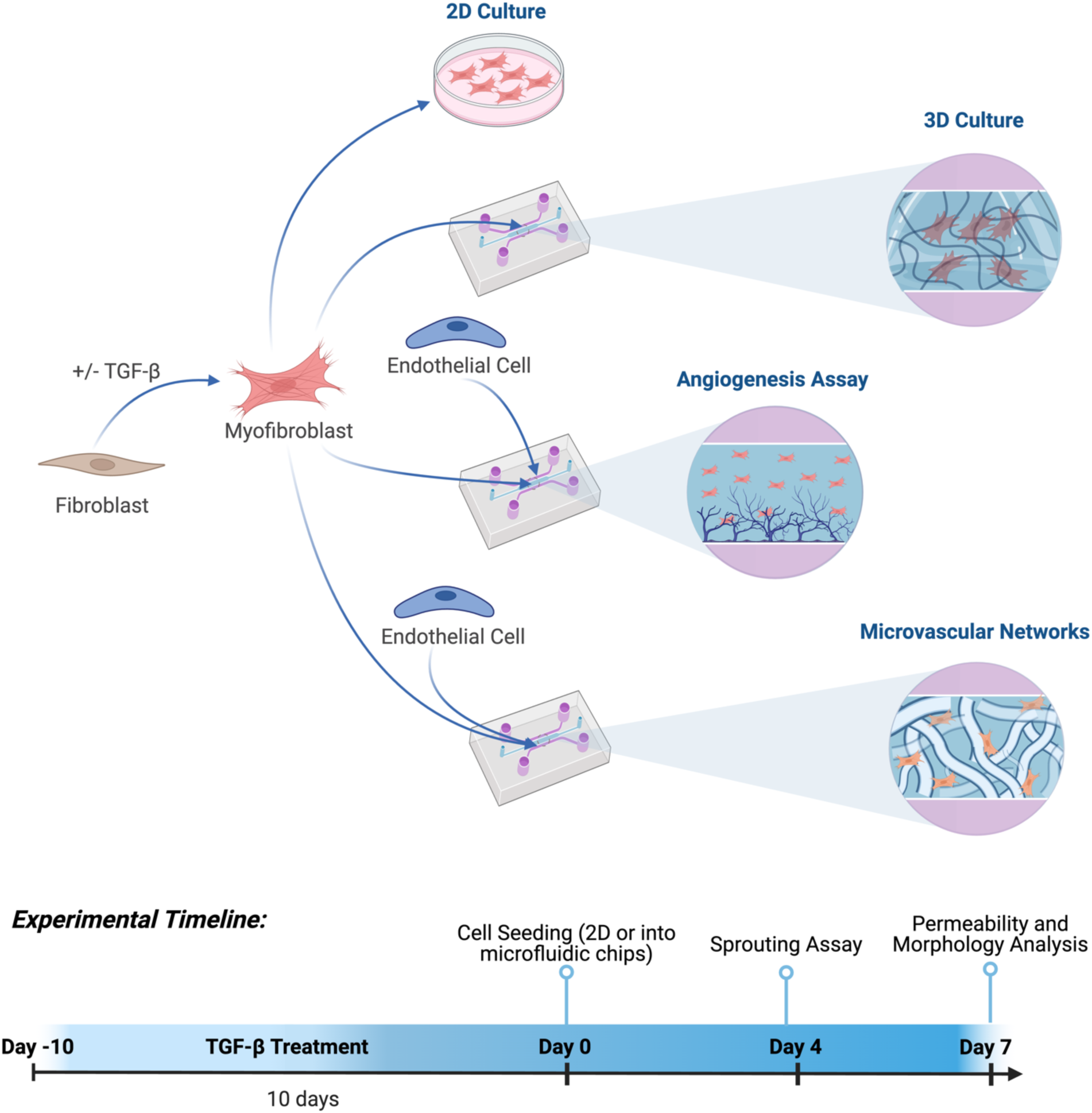
Experimental platforms and timeline. Scheme illustrating culture of fibroblasts or myofibroblasts on a 2D surface or in 3D, suspended in a hydrogel within microfluidic chips. Endothelial cells were co-cultured with fibroblasts or myofibroblasts in 3D to study angiogenesis and vasculogenesis of microvascular networks. The experimental timeline indicates the duration and timing of cell culture, TGF-β treatment, and assays.

**Fig. 2.**
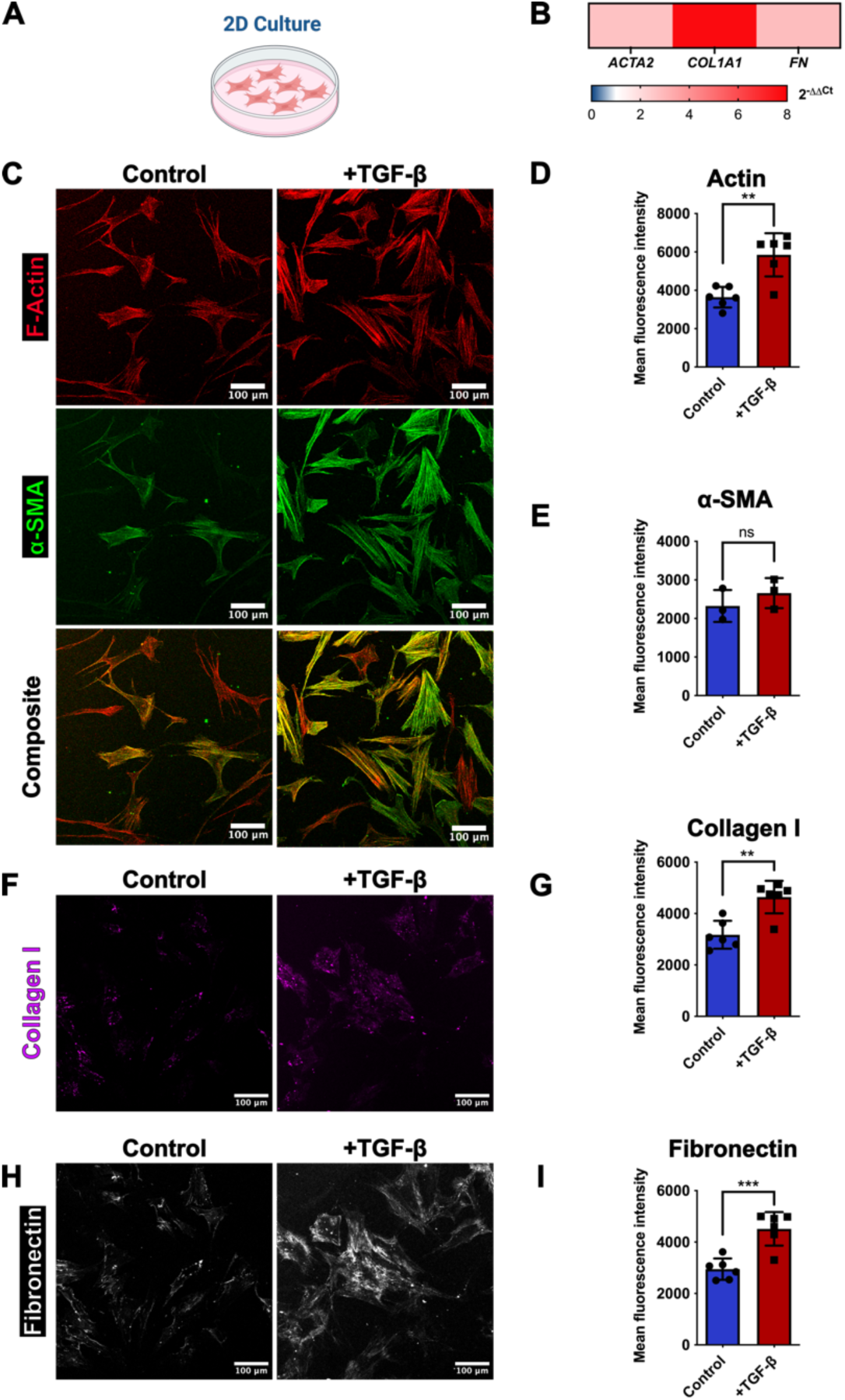
Generation of myofibroblasts in 2D. (**A**) Fibroblasts were cultured in 2D and treated with TGF-β for 10 days. (**B**) Expression of gene transcripts in cells after treatment with TGF-β for 10 days. Expression levels are relative to untreated controls. n=3 independent plates for each group. (**C**) Representative images of F-actin (red) and α-SMA (green) in control and TGF-β-treated cells. Quantification of mean fluorescence (**D**) actin (P=0.002) and (**E**) α-SMA signal (P=0.17) in control and TGF-β-treated cells. (**F**) Representative images of collagen I (magenta) in control and TGF-β-treated cells. (**G**) Quantification of mean fluorescence collagen I signal (P=0.002). (**H**) Representative images of fibronectin (white) in control and TGF-β-treated cells. (**I**) Quantification of mean fluorescence fibronectin signal (P=0.0006). n=3-6 coverslips from 3 independent experiments for each group in D, E, G, and I. Reported P values are based on unpaired t-tests.

### The lung myofibroblast phenotype is maintained in 3D after removal of TGF-β

One method of generating engineered microvascular networks *in vitro* is through self-assembly of human endothelial cells and fibroblasts embedded in a fibrin gel in a microfluidic device (*17*, *18*). This process takes approximately 4-7 days of co-culture, during which time the endothelial cells migrate, connect, and organize into lumenized vessel structures. Given that the objective of this study is to isolate the effect of myofibroblasts on vascularization, it is important to avoid treating endothelial cells with exogenous TGF-β to isolate the effect of the myofibroblasts. Therefore, we chose to incorporate pre-differentiated lung myofibroblasts instead of differentiating them *in situ*. It was also critical to verify that the myofibroblast phenotype was maintained in the 3D cell culture conditions used in self-assembled microvascular networks. As previously established, lung fibroblasts were grown for 10 days in 2D without TGF-β (referred to as “fibroblasts” in all subsequent experiments) or with TGF-β (referred to as “myofibroblasts” in all subsequent experiments) (**Fig. 1**). Fibroblasts or myofibroblasts were detached and embedded in fibrin in a microfluidic device and subsequently cultured in endothelial medium for 7 days to mimic the conditions employed in vascular self-assembly, but lacking the endothelial cells necessary for the formation of vasculature (**Fig. 1**, **Fig. 3A**). After 7 days of culture, gene and protein expression of α-SMA, collagen I, and fibronectin were assessed with RT-qPCR and immunofluorescence. Increased mRNA levels of ACTA2 (2.3-fold), COL1A1 (2.3-fold), and FN1 (1.3-fold) were maintained in myofibroblasts relative to fibroblasts (**Fig. 3B**), similar to what was observed at the end of 2D culture (**Fig. 2B**). At the protein level, myofibroblasts showed higher fluorescence intensity for actin (**Fig. 3C, D**) and α-SMA (**Fig. 3C, E**) compared to fibroblasts. Although not a statistically significant difference, the fluorescence intensity for collagen I and fibronectin was also increased in myofibroblasts (**Fig. 3F-I**). Overall, these results show that the myofibroblast phenotype was maintained in 3D endothelial cell-compatible culture for 7 days despite the withdrawal of TGF-β prior to incorporation into the 3D system.

**Fig. 3.**
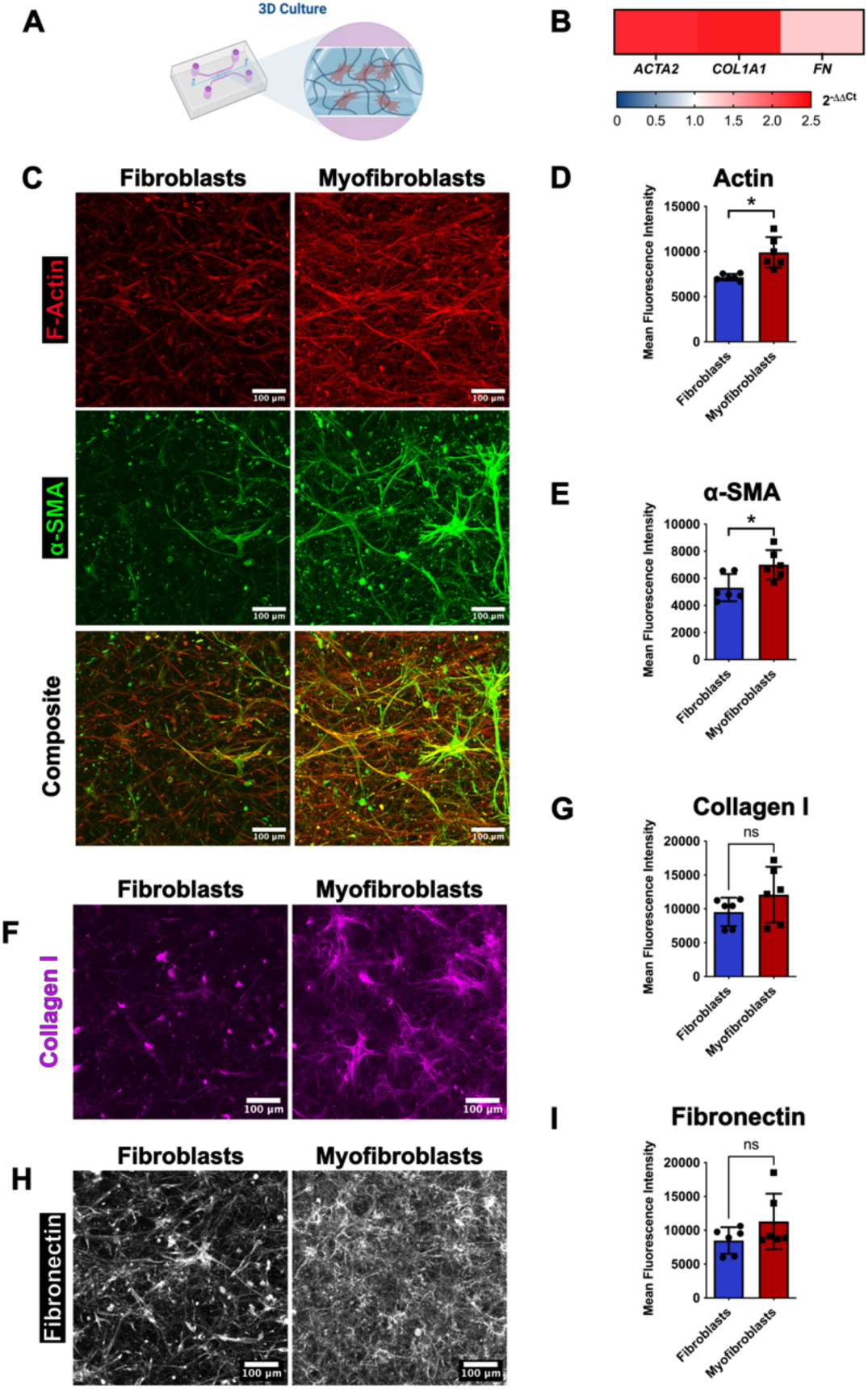
Maintenance of myofibroblast phenotype in 3D culture. (**A**) Fibroblasts or myofibroblasts were cultured in 3D within fibrin gels housed in microfluidic devices. (**B**) Expression of gene transcripts in fibroblasts or myofibroblasts after 7 days of culture in 3D. Expression levels are relative to fibroblasts. n=6 pooled devices for each group. (**C**) Representative images of F-actin (red) and α-SMA (green) in fibroblasts and myofibroblasts. Quantification of mean fluorescence (**D**) actin (P=0.003) and (**E**) α-SMA (P=0.02) signal in fibroblasts and myofibroblasts. (**F**) Representative images of collagen I in fibroblasts and myofibroblasts. (**G**) Quantification of mean fluorescence collagen I signal (P=0.20). (**H**) Representative images of fibronectin (white) in fibroblasts and myofibroblasts (**I**) Quantification of mean fluorescence fibronectin signal (P=0.16). n=6 devices from 3 independent experiments for each group in D, E, G, and I. Reported P values are based on unpaired t-tests.

### Myofibroblasts suppress endothelial cell angiogenic sprouting

We next assessed the interaction of fibroblasts or myofibroblasts with human endothelial cells by investigating their influence on sprouting angiogenesis. To this end, lung fibroblasts or myofibroblasts were embedded in fibrin in the central gel channel of the microfluidic device and fluorescent endothelial cells were applied as a monolayer on one vertical face of the gel channel (**Fig. 1**, **Fig. 4A**). Endothelial cell sprouting was assessed after 4 days using confocal microscopy (**Fig. 1**). Image analysis revealed that fibroblasts induced significantly more angiogenesis compared to myofibroblasts, quantified by the coverage by endothelial cells (sprouting area) (**Fig. 4B,C**), demonstrating that myofibroblasts are less pro-angiogenic.

**Fig. 4.**
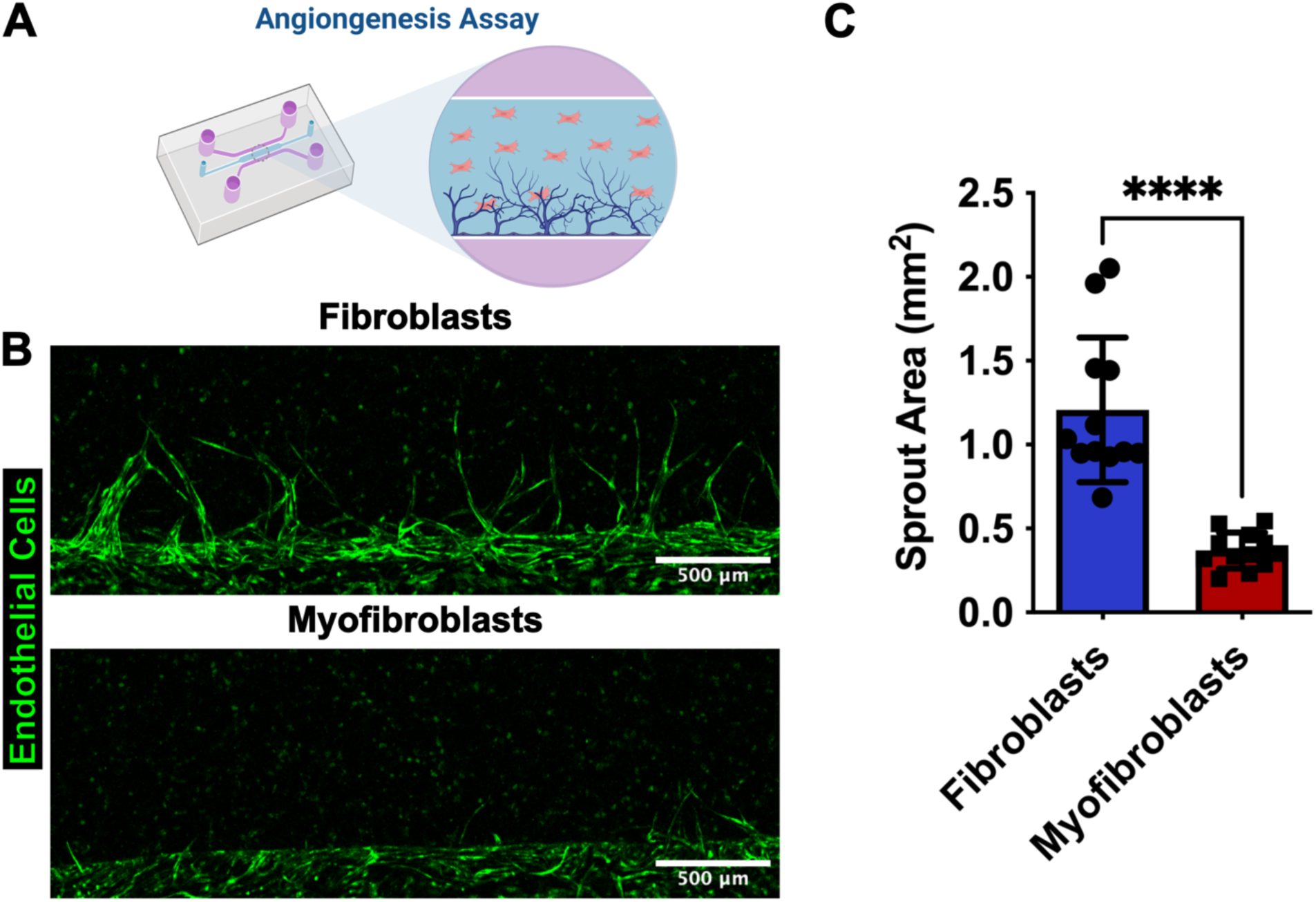
Effects of myofibroblasts on angiogenic sprouting. (**A**) Endothelial cells were co-cultured with either fibroblasts or myofibroblasts in 3D for angiogenesis analysis. (**B**) Representative images of endothelial cell sprouts (green) co-cultured with fibroblasts or myofibroblasts. (**C**) Endothelial sprout area in cultures with fibroblasts or myofibroblasts (P<0.0001, unpaired t-test). n=12 devices from 3 independent experiments for each group.

### Myofibroblasts reduce the diameter and increase the permeability of vessels in engineered microvascular networks

In the next step, we assessed the effects of myofibroblasts on vasculogenesis and vascular barrier function. Microvascular networks were formed by self-assembly of human endothelial cells in combination with either lung fibroblasts or myofibroblasts (**Fig. 1**, **Fig. 5A**). Seven days after seeding, microvascular networks were perfused with fluorescent dextran (70 kDa) and imaged with a confocal microscope to assess vessel morphology and permeability (**Fig. 1**). Myofibroblasts generated microvascular networks with thinner vessels compared to fibroblasts (**Fig. 5B**). Morphology analysis revealed that microvascular networks containing myofibroblasts and fibroblasts had a similar number of branches (**Fig 5C**) and total vessel length (**Fig. 5D**). However, microvascular networks with myofibroblasts displayed a significantly smaller vessel diameter (**Fig. 5E**) and a higher permeability to 70 kDa dextran (**Fig. 5F**), suggesting that they have impaired barrier function compared to vasculature generated in co-culture with fibroblasts.

**Fig. 5.**
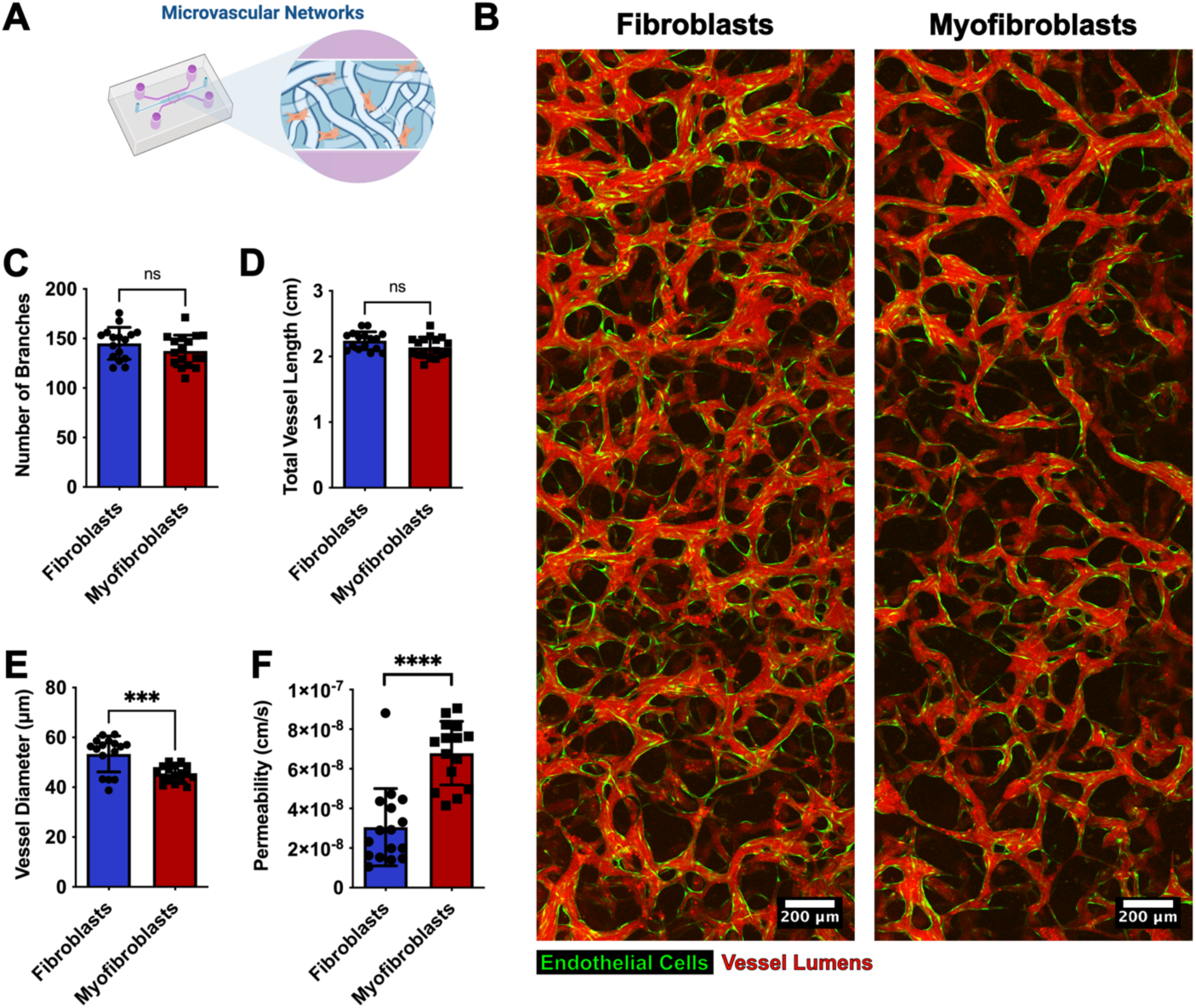
Effects of myofibroblasts on vasculogenesis. (**A**) Microvascular networks were formed with endothelial cells and either fibroblasts or myofibroblasts. (**B**) Representative images of microvascular networks (endothelial cells in green) perfused with fluorescent dextran to indicate vessel lumens (red) in samples with fibroblasts or myofibroblasts. Quantification of (**C**) number of vessel branches (P=0.18), (**D**) total length of vasculature (P=0.06), (**E**) average vessel diameter (P=0.0002), and (**F**) vascular permeability (P<0.0001, Mann-Whitney U test) of microvascular networks with fibroblasts or myofibroblasts. n=16 devices from 3 independent experiments in each group. Reported P values for all measurements other than permeability are based on unpaired t-tests.

### Myofibroblasts secrete high levels of TGF-β1 and low levels of VEGF

In order to understand the mechanisms responsible for the anti-angiogenic, anti-vasculogenic, and pro-fibrotic activities of myofibroblasts, we profiled the secreted factors related to angiogenesis and fibrosis in supernatant collected and pooled over the 7 days of 3D monoculture in the microfluidic devices. Among the cytokines that were highly secreted, we found a 1.36-fold increase in TGF-β1 in supernatant from devices with myofibroblasts versus those with fibroblasts (**Fig. 6A**), as well as a 0.53-fold change in vascular endothelial growth factor-A (VEGF) (**Fig. 6B**). We also measured the same cytokine panel in supernatant from microvascular networks containing either fibroblasts or myofibroblasts to assess the crosstalk between endothelial cells and lung (myo)fibroblasts (**Fig. 6C**). We found that among the highly secreted cytokines, several were differentially expressed in vasculature with myofibroblasts relative to those with fibroblasts, including granulocyte-colony stimulating factor (G-CSF, 1.64-fold), VEGF (1.30-fold), cytokine C-C motif ligand 2 (CCL2, 1.56-fold), and interleukin-8 (IL-8, 1.90-fold) (**Fig. 6C**). These results highlight several potential mechanisms by which myofibroblasts influence angiogenesis and vasculogenesis in the microphysiological model of lung fibrosis.

**Fig. 6.**
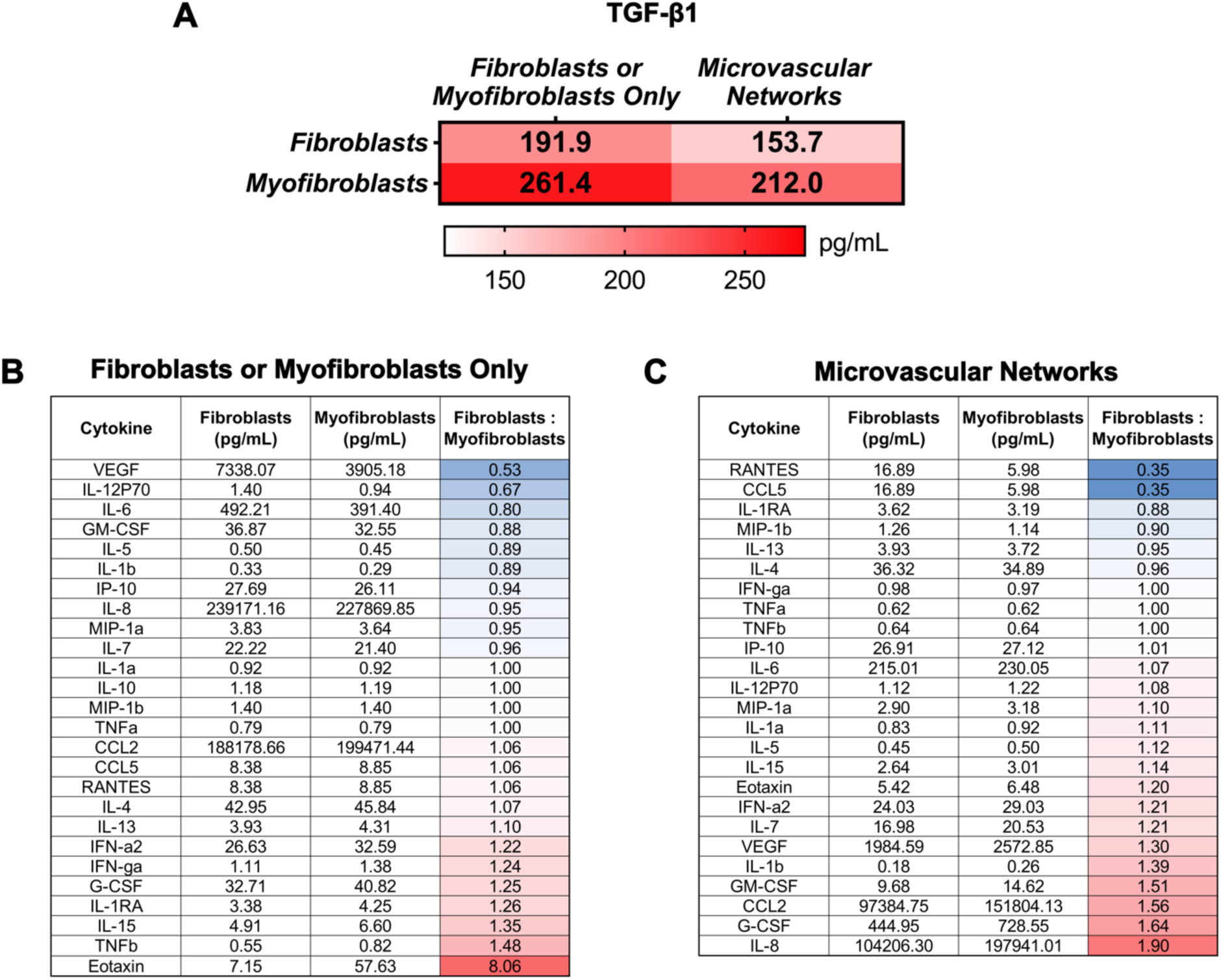
Cytokines secreted in fibrosis model. (**A**) TGF-β1 secreted in 3D monocultures of fibroblasts or myofibroblasts or in microvascular networks containing either fibroblasts or myofibroblasts. (**B**) Panel of cytokines secreted in 3D monocultures of fibroblasts or myofibroblasts. (**C**) Panel of cytokines secreted by microvascular networks containing either fibroblasts or myofibroblasts cultured in 3D. n=6 pooled devices per group.

### Blocking TGF-β receptor or supplementing VEGF rescues vessel morphology and permeability

Finally, we modulated the concentration or function of the identified cytokine targets to determine if this could reverse the effects of myofibroblasts on microvascular structure and function. We focused on IL-8 due to its high concentration and differential secretion between microvascular networks containing fibroblasts and myofibroblasts. Based on the data from 3D monocultures, we also examined TGF-β1 and VEGF. Repeating the methods from the vasculogenesis experiments, lung fibroblasts or myofibroblasts were seeded with endothelial cells in fibrin in the microfluidic device to form microvascular networks. However, in this iteration, devices with myofibroblasts were instead cultured in endothelial medium supplemented with one of the following: vehicle (dimethylsulfoxide), an anti-IL-8 blocking antibody, the TGF-β type I receptor/ALK5 inhibitor SB-431542, or human recombinant VEGF. After 7 days of culture, microvascular networks were perfused with dextran and imaged to compare vessel morphology and permeability across treatment conditions (**Fig. 7A**). In the vehicle controls, microvascular networks containing myofibroblasts showed a reduced number of branches (**Fig. 7B**), lower total vessel length (**Fig. 7C**), decreased vessel diameter (**Fig. 7D**), and increased vessel permeability (**Fig. 7E**) compared to microvascular networks with fibroblasts. While the anti-IL-8 blocking antibody did not result in any significant effects on vessel morphology and permeability (**Fig. 7B-E**), SB-431542 restored total vessel length (**Fig. 7C**), vessel diameter (**Fig. 7D**), and permeability (**Fig. 7E**) to levels similar to those of microvascular networks containing fibroblasts. Similarly, supplementation with recombinant VEGF significantly increased the number of branches (**Fig. 7B**) and total vessel length (**Fig. 7C**), but not vessel diameter (**Fig. 7D**). Moreover, VEGF restored the endothelial barrier function of the microvascular networks containing myofibroblasts by reducing vessel permeability (**Fig. 7E**). Together, these results show that myofibroblasts influence vessel morphology and permeability via the increased secretion of TGF-β1 and reduced secretion of VEGF, while the secretion of IL-8 may not play a role in these phenomena. Overall, the rescue of the vessel morphology and permeability to values resembling those of normal vasculature by SB-431542 and VEGF showcases the ability of our microphysiological model of lung fibrosis to screen anti-fibrotic drugs targeting vascular structure and function.

**Fig. 7.**
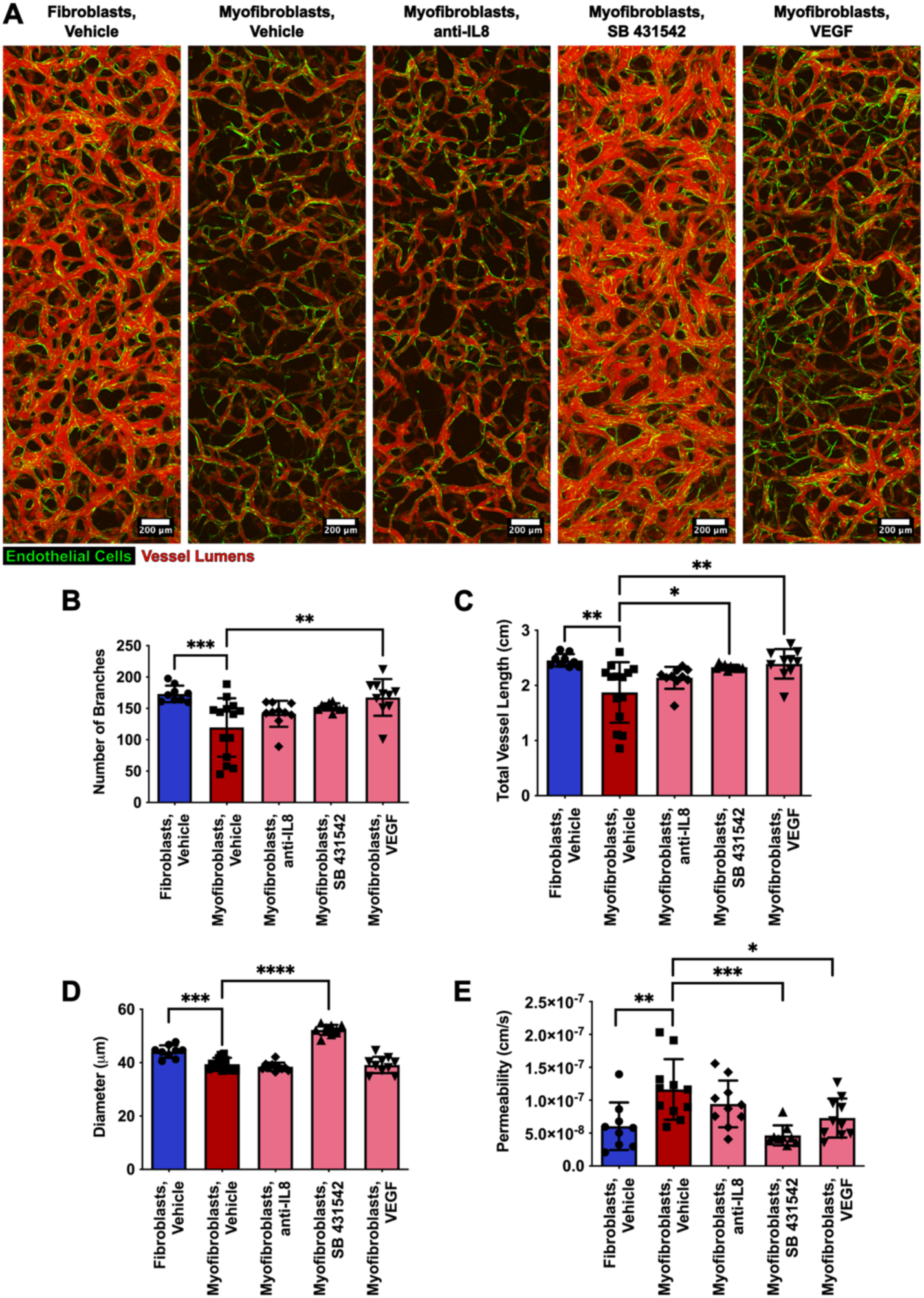
Effects of select compounds on microvascular structure and function. (**A**) Representative images of microvascular networks (endothelial cells in green) perfused with fluorescent dextran to indicate vessel lumens (red) in samples with fibroblasts or myofibroblasts and treated with vehicle or select compounds. Quantification of (**B**) number of vessel branches (P=0.0009 for Fibroblasts, Vehicle vs. Myofibroblasts, Vehicle; P=0.002 for Myofibroblasts, Vehicle vs. Myofibroblasts, VEGF), (**C**) total length of vasculature, (P=0.001 for Fibroblasts, Vehicle vs. Myofibroblasts, Vehicle; P=0.01 for Myofibroblasts, Vehicle vs. Myofibroblasts, SB 431542; P=0.003 for Myofibroblasts, Vehicle vs. Myofibroblasts, VEGF) (**D**) average vessel diameter (P=0.0002 for Fibroblasts, Vehicle vs. Myofibroblasts, Vehicle; P<0.0001 for Myofibroblasts, Vehicle vs. Myofibroblasts, SB 431542), and (**E**) vascular permeability (P=0.006 for Fibroblasts, Vehicle vs. Myofibroblasts, Vehicle; P=0.0004 for Myofibroblasts, Vehicle vs. Myofibroblasts, SB 431542; P=0.04 for Myofibroblasts, Vehicle vs. Myofibroblasts, VEGF) of microvascular networks with fibroblasts or myofibroblasts and treated with vehicle or anti-fibrotic drug. n=9-11 devices from 2 independent experiments in each group. Reported P values are based on one-way ANOVA tests with multiple comparisons.

## Discussion

The goal of this study was to develop a pathophysiologically relevant model of microvasculature in lung fibrosis to investigate the crosstalk between myofibroblasts and endothelial cells. In a first step, we showed that human lung fibroblasts cultured in 2D differentiate into myofibroblasts after treatment with 1 ng/ml TGF-β for 10 days, as measured by increased expression of actin and α-SMA and the increased production of collagen I and fibronectin (**Fig. 2**). These results are in line with results obtained with similar protocols aimed at differentiating fibroblasts into myofibroblasts *in vitro* (*11*, *16*, *19–22*). Importantly, the changes observed upon treatment with TGF-β (cell morphology, α-SMA expression, and extracellular matrix production) recapitulate the characteristics of lung myofibroblasts observed in rat models of lung fibrosis (*23*, *24*) and patients with lung fibrosis (*25*).

It was crucial to verify that myofibroblasts maintained their phenotype once TGF-β was withdrawn and they were introduced into a 3D hydrogel in order to avoid treating microvascular networks with exogenous TGF-β. This enabled us to build a more accurate model of the pathophysiology of lung fibrosis in which all the paracrine signaling originates from cellular sources rather than exogenous supplementation. This feature is notably difficult to achieve, as myofibroblast differentiation depends on TGF-β signaling, adhesion signals, and integrin signaling (*19*). Here, we report a myofibroblast phenotype that remains stable in 3D for at least 7 days in absence of TGF-β supplementation (**Fig. 3**). We attribute this result to a long pre-treatment duration of 10 days, because we observed in an optimization phase that the myofibroblast phenotype was not maintained in 3D after pre-conditioning fibroblasts with 1 ng/ml and higher concentrations of TGF-β for only 2 days (data not shown). We speculate that a longer pre-treatment induces a reprogramming of the fibroblasts, which then commit to a more stable myofibroblast phenotype. In the next step, we used 3D microphysiological systems of increasing complexity to perform functional assays and study the influence of myofibroblasts on angiogenesis, vasculogenesis, and permeability (**Fig. 1**). We show that myofibroblasts cultured in 3D impaired angiogenesis compared to fibroblasts, as measured by the coverage area formed by endothelial sprouts after the cells were seeded on the side of the gel and allowed to grow into the 3D matrix containing fibroblasts or myofibroblasts (**Fig. 4**). These results suggest that myofibroblasts have an anti-angiogenic activity, corroborating clinical studies reporting decreased abundance of alveolar capillaries and elevated expression of anti-angiogenic factors in lung tissue and serum in patients with IPF (*8*, *25–27*). In particular, this model is more representative of extensive fibrotic lesions, which are characterized by decreased microvascular density relative to control lungs, whereas regions of minimal fibrosis display higher microvascular density (*8*, *25*).

We also show that the presence of myofibroblasts impacts vasculogenesis, as myofibroblasts co-cultured with endothelial cells resulted in microvasculature with narrower vessels and increased permeability compared to models containing normal fibroblasts (**Fig. 5**). These results are in agreement with studies reporting reduced vascular area, in addition to reduced vascular density, in biopsies of lung fibrosis patients compared to control biopsies (*28*). Moreover, vascular regression (decreased luminal area and perimeter of vessels) was shown to increase with increasing levels of lung fibrosis (*29*). The permeability results are also consistent with observations of enhanced and persistent vascular permeability characteristic of the fibrotic lung (*30*).

We hypothesized that the anti-angiogenic, anti-vasculogenic, and pro-fibrotic activities of myofibroblasts in our models were due to paracrine signaling, given that fibroblasts or myofibroblasts are not in direct physical contact with endothelial cells in our angiogenesis assay until endothelial cells invade the 3D matrix. Through the analysis of the secretome, we determined that myofibroblasts in 3D monoculture have an increased secretion of TGF-β1 and a reduced secretion of VEGF compared to normal fibroblasts (**Fig. 6A,B**). These results suggest that myofibroblasts produce TGF-β1 even after the withdrawal of exogenous sources, corroborating a stable reprogramming of fibroblasts into myofibroblasts through the persistence of autocrine TGF-β1 signaling. Moreover, the considerably diminished secretion of VEGF by myofibroblasts is a likely cause of decreased angiogenesis and vasculogenesis, as VEGF is a crucial growth factor for these processes. This result aligns with a study showing that reduced vascular density in lung fibroblastic foci is accompanied by reduced VEGF concentration and increased TGF-β1 concentration (*27*). Reduced levels of VEGF were also observed in bronchoalveolar lavage fluid of patients with lung fibrosis (*31*, *32*). Interestingly, while TGF-β1 secretion was consistently higher in the 3D culture with myofibroblasts (alone or in co-culture with endothelial cells), VEGF secretion was not consistent between myofibroblast monoculture and co-culture. In this case, VEGF concentrations were significantly lower in devices with myofibroblasts alone (**Fig. 6B**) but elevated in microvascular networks containing myofibroblasts (**Fig. 6C**). We speculate that this is due to increased VEGF secretion by the endothelial cells in the fibrotic model (*33–35*) and/or their impaired capacity to bind VEGF (*34*). The increased secretion of the G-CSF, CCL2, and IL-8 cytokines in the microvascular networks containing myofibroblasts (**Fig. 6C**) further suggests that our model recapitulates the fibrotic lung microenvironment, as these cytokines are present in high levels in IPF (*36–39*) and promote lung fibrosis by attracting immune cells to the injured lung tissue, which then contribute to inflammation (*36–40*).

Focusing on the modulation of selected cytokines (IL-8, TGF-β1, and VEGF), we show that the effects of the myofibroblasts on the microvasculature can be rescued through targeted pharmacological interventions. While IL-8 inhibition was ineffective at reverting vascular morphological changes and permeability to normal conditions, either inhibition of TGF-β type I receptor/ALK5 with SB-431542 or supplementation with recombinant VEGF successfully rescued vascular morphology and barrier function, with SB-431542 displaying the strongest effect (**Fig. 7**). These findings are in line with studies showing that ALK5 inhibition protects endothelial cells from apoptosis (*41*), enhances their growth and integrity (*42*), and reduces vascular permeability (*43*). Moreover, VEGF delivery reduced endothelial cell apoptosis and increased vascularization in a rat model of IPF induced by adenoviral delivery of active TGF-β1 (*34*). Therefore, this study validates the capacity of this vascular model to screen therapeutic approaches to treat lung fibrosis. It is important to acknowledge that while these microvascular models recapitulate key aspects of lung fibrosis, they do not fully recapitulate the spatial and temporal heterogeneity present in fibrotic lung tissue (*42*). However, additional aspects of disease could be incorporated by integrating cells collected from different regions of a fibrotic lung or harvested at varying stages of disease. To improve physiological relevance, future models could also include lung-specific endothelial cells, such as primary lung endothelial cells or induced pluripotent stem cell-derived endothelial cells, and lung-specific mechanical stimuli, such as continuous vascular flow and breathing-induced stretching.

The effect of myofibroblasts on vasculature in lung fibrosis is poorly understood, as evidenced by controversial scientific results. This highlights the need for advanced microphysiological systems that recapitulate the pathophysiology of lung fibrosis and allow to dissect the effect of myofibroblasts in co-culture with endothelial cells. The microvascular model presented here captures key aspects of lung fibrosis, including the presence of lung myofibroblasts that maintain their characteristic phenotype in 3D for at least 7 days in absence of exogenous TGF-β. Compared to fibroblasts, myofibroblasts exerted anti-angiogenic and anti-vasculogenic effects on endothelial cells, as measured by reduced endothelial sprouting, altered vascular morphology, and increased vascular permeability. Myofibroblasts secreted increased levels of TGF-β1 and decreased levels of VEGF, and targeted pharmacological interventions including TGF-β type I receptor/ALK5 inhibition with SB-431542 and supplementation with recombinant VEGF rescued vessel morphology and permeability. As such, this model holds great promise as a translational assay to screen anti-fibrotic drugs and drugs aimed at vascular normalization in lung fibrosis.

## Materials and methods

### Myofibroblast induction

Primary human lung fibroblasts (Lonza, CC-2512, passage 8) were thawed and seeded in two 75 cm^2^ flasks in Fibrolife S2 medium (Lifeline Cell Technology, LL-0011). After 24 hours of recovery from cryopreservation, the media was replaced with DMEM/F12 (Gibco 11320033) supplemented with 1% penicillin/streptomycin (Gibco, 15140122) and 0.5% fetal bovine serum (Gibco, 26140079). One flask was supplemented with 1 ng/ml TGF-β1 (PeproTech, 100-21-10UG) to induce the myofibroblast phenotype. Fibroblasts (with and without TGF-β) were cultured for 10 days with media changes every 2 days before harvesting for immunofluorescent staining, angiogenic sprouting assays, or vasculogenesis experiments.

### Microfluidic device fabrication

Molds of microfluidic devices with a central gel channel measuring 3×10×0.5 mm were designed in Fusion 360 (Autodesk) and milled using a Computer Numerical Control (CNC) Milling Machine (Bantam Tools). Devices were subsequently fabricated as previously described (*44*, *45*). Briefly, polydimethylsiloxane (PDMS, Dow Corning Sylgard 184, Ellsworth Adhesives) mixed at a ratio of 10:1 was degassed for 30 minutes, poured into molds, degassed a second time for 30 minutes, and cured in an oven at 60°C overnight. Cured PDMS devices were trimmed and ports were punched out using biopsy punches (Integra Miltex) prior to autoclaving. Individual devices and clean glass coverslips (#1, VWR) were plasma treated (Expanded Plasma Cleaner, Harrick Plasma), bonded, and placed in an oven at 70°C overnight prior to use.

### RNA extraction and RT-qPCR analysis of 2D and 3D samples

RNA was extracted with the RNeasy Mini Kit (Qiagen) according to the manufacturer’s recommendations. Briefly, cells in 2D were lysed in RLT buffer with beta-mercaptoethanol and scraped from the culture surface. RNA from cells in 3D was extracted by first removing the gel from a microfluidic device after cutting a window in the PDMS. The matrix was dissociated in 100 μg/ml Liberase TM (Millipore Sigma) until a single cell suspension was achieved, and cells were washed and pelleted before lysis and RNA extraction. cDNA was generated using SuperScript III First-Strand Synthesis SuperMix for qRT-PCR (Thermofisher). RT-qPCR was completed with the Applied Biosystems 7300 Fast real-time PCR system and software using Power SYBR Green PCR Master Mix (Thermofisher). Primer sequences were selected from the Massachusetts General Hospital Primer Bank and synthesized by Integrated DNA Technologies (TGFB1: forward primer GGCCAGATCCTGTCCAAGC, reverse primer GTGGGTTTCCACCATTAGCAC); ACTA2: forward primer AAAAGACAGCTACGTGGGTGA, reverse primer GCCATGTTCTATCGGGTACTTC; COL1A1: forward primer GAGGGCCAAGACGAAGACATC, reverse primer CAGATCACGTCATCGCACAAC; FN1: forward primer CGGTGGCTGTCAGTCAAAG, reverse primer AAACCTCGGCTTCCTCCATAA). Fold-change was computed using the 2^-ΔΔCT^ method against the h36B4 housekeeping gene (*46*).

### Immunofluorescence staining of 2D & 3D samples and image analysis

The samples were fixed with 4% paraformaldehyde (Electron Microscopy Sciences, 50-259-96) for 15 minutes and washed twice with PBS (Gibco, 10010049). For intracellular imaging, the samples were permeabilized with 0.1% Triton X-100 (Invitrogen, HFH10) for 5 minutes at room temperature and then washed with PBS twice. Subsequently, all samples were treated with 5% goat serum (Invitrogen, 16210072) for 45 minutes at room temperature. After two additional washes with PBS, the samples were incubated with antibodies recognizing α-SMA (Cell Signaling Technology, 19245, diluted 1:200 in PBS for intracellular imaging), collagen 1, (Cell Signaling Technology, 39952, diluted 1:200 in PBS for extracellular imaging), and fibronectin (R&D Systems, MAB1918, diluted 1:200 in PBS for extracellular imaging) overnight at 4°C. On the following day, the samples were washed twice with PBS and then incubated with secondary antibodies diluted 1:200 in PBS (Alexa Fluor 488-Donkey anti-Rabbit, Invitrogen, A-21206, for intracellular imaging, and Alexa Fluor 488-Goat anti-mouse, Invitrogen, A-11001 for extracellular imaging) along with stains for F-actin (Alexa Fluor 594-Phalloidin, Invitrogen, A12381, 1:200 dilution in PBS) and nuclei (4’,6-diamidino-2-phenylindole, Sigma-Aldrich, D8417, diluted 1:10000 in PBS), for 1 hour at room temperature. Finally, the samples were washed twice with PBS and stored in PBS until the day of imaging. Imaging was performed at 20x using an Olympus FLUOVIEW FV1200 confocal laser scanning microscope. Images were analyzed using ImageJ (NIH, U.S.A) by summing confocal slices and computing image intensity statistics.

### Angiogenic sprouting assay

Pooled human umbilical vein endothelial cells (HUVECs, Lonza, C2519A, passage 7) transduced to express cytoplasmic green fluorescent protein (*47*) were cultured in 75 cm^2^ flasks in VascuLife VEGF medium (Lifeline Cell Technology, LL-0003) with all supplements in the bullet kit, with the exception of heparin sulfate (only 25% of the recommended concentration was used) prior to use in microfluidic devices.

After detachment from cell culture flasks, fibroblasts or myofibroblasts were resuspended in cold VascuLife medium containing 3.6 U/ml thrombin (Sigma-Aldrich, T4648), mixed with an equal volume of 5 mg/mL fibrinogen (Sigma-Aldrich, F8630), and seeded in the central channel of microfluidic devices at final density of 1.5×10^6^ cells/ml. The cell-laden gel solution was allowed to polymerize at 37° C for approximately 15 minutes before warm VascuLife media was added to the side channels. After 24 hours, a solution of fibronectin (30 μg/ml, Millipore Sigma, FC010) was added to a side channel and incubated for 30 minutes. Then, the fibronectin solution was aspirated and 40 μl of a suspension of HUVECs (1×10^6^ cells/ml) were added to the channel. The device was rotated 90° for 15 minutes to permit cell settling on the surface of the fibrin channel. The devices were then righted and incubated for an additional 15 minutes to permit cell adhesion before replacing the medium with fresh VascuLife. Medium was changed daily for 4 days, at which time the devices were imaged with a confocal microscope (Olympus FV1000) at 37°C and using a 4x objective to identify sprouting of the fluorescent endothelial cells into the fibroblast or myofibroblast-laden gel compartment. Confocal z-stack images were acquired at a resolution of 0.25 pixels/μm at a z-spacing of 27 μm. Maximum intensity projections of image z-stacks were cropped to exclude any signal from endothelial cells growing in the media channel. Then, a global threshold was applied for binarization of the sprouts and quantification of the area occupied by sprouts was performed with ImageJ.

### Formation of microvasculature (vasculogenesis assay)

After detachment from cell culture flasks, HUVECs (same as above) and fibroblasts or myofibroblasts were resuspended in cold VascuLife medium containing 3.6 U/ml thrombin (Sigma-Aldrich, T4648), mixed with an equal volume of 5 mg/mL fibrinogen (Sigma-Aldrich, F8630), and seeded in the central channel of microfluidic devices at final density of 7×10^6^ HUVECs/ml and 1.5×10^6^ (myo)fibroblasts/ml. The cell-laden gel solution was allowed to polymerize at 37° C for approximately 15 minutes before warm VascuLife media was added to the side channels. Medium was changed daily, and starting on day 5 of culture, 1 ml syringes (plungers removed and cut in height) were placed in the media ports on one of the media channels of each device. The two syringes were filled with 200 μl of medium each to produce flow across the gel channel to the open medium channel on the opposite side. Microvascular network morphology and permeability was quantified on day 7 of 3D co-culture.

### Analysis of vascular permeability

A 0.1 mg/ml solution of fluorescently labeled dextran (Texas-Red, 70 kDa, Invitrogen, D1864) was introduced into the microvascular networks, and the devices were imaged at 37°C using a confocal microscope (Olympus FV1000, 10x objective). Two sets of confocal z-stack images, spanning a total height of 200 μm, were acquired at a resolution of 0.5 pixels/μm at a z-spacing of 5 μm for each region of interest, 12 minutes apart. Analysis of the changes in the intensity of the fluorescence signal in the vessels and the matrix over time was performed to quantify vascular permeability as previously described (*45*).

### Analysis of vascular morphology

Maximum intensity projection images were generated from confocal z-stacks of vessels perfused with dextran that were previously acquired for permeability analysis. AutoTube software (*48*) was used to calculate total vascular length, number of branches, and vascular diameter. Images were processed by applying adaptive histogram equalization, illumination correction, and denoising using block-matching and 3D filtering. Thresholding to detect tubes was performed using the Multi-Otsu method and regions measuring less than 1% of the image were removed. Additionally, short ramifications (less than 30 pixels) were removed and branch points with a spatial distance less than 15 pixels were merged prior to vessel analysis.

### Analysis of secreted cytokine (ELISA and multiplex cytokine assay)

Medium in microfluidic devices was changed daily in devices with (myo)fibroblasts monoculture or co-culture with endothelial cells. The supernatant was collected daily prior to the addition of fresh medium and pooled from all devices in the same group. Furthermore, the supernatant samples collected during the 7 days of culture were combined to represent average cytokine concentrations across a week of culture. The concentration of TGF-β1 was quantified using a Quantikine ELISA kit (R&D Systems, Human/Mouse/Rat/Porcine/Canine TGF-β1, D8100C) according to the manufacturer’s recommendations. Absorbance at 450 nm was detected with a SpectraMax i3x microplate reader (Molecular Devices), with correction at 540 and 570 nm. The same supernatant was also analyzed with a multiplexed cytokine assay (Human cytokine/chemokine magnetic bead panel, Millipore Sigma, HCYTMAG-60K0PX30) on a Luminex Magpix system (Millipore Sigma) to determine concentrations of secreted proteins using five-parameter curve-fitting models.

### Interventions to rescue vascular structure and function

The effect of several compounds on vasculogenesis was tested on microvascular networks with fibroblasts or myofibroblasts that were prepared as described above. Starting on the day of seeding into microfluidic devices, vehicle-control microvascular networks were cultured in VascuLife media containing 1:2000 dimethylsulfoxide (ATCC, 4-X), while treated microvascular networks were cultured in VascuLife with 20 μg/ml human IL-8/CXCL8 antibody (R&D Systems, MAB208), 10 μM SB 431542 (Tocris, 1614/1), or 0.05 μg/ml human VEGF (PeproTech, 100-20-10UG). On day 7 of culture, devices were imaged, and permeability and vascular morphology was analyzed as detailed above.

#### Statistics

Data were checked for normality and presented as mean ± standard deviation. The data points provided in the graphs represent single devices or coverslips from two or three independent experiments. Statistical analysis of the data was performed with the software GraphPad Prism. The specific statistical test used is reported in each figure caption. Mean differences with p value < 0.05 were considered statistically significant.

## Author contributions

Conceptualization: EC, AB, ECK, SES

Methodology: EC, AB, ECK, SES

Investigation: EC, AB, ECK, TT, SD, SES,

Visualization: AB

Supervision: DAB, SES, RDK

Writing original draft: EC, AB, SES

Writing review & editing: EC, AB, ECK, TT, SD, DAB, SES, RDK

## Data and materials availability

All data are available in the main text.

